# P2X receptors in *Aplysia californica*: Chemosensory systems, bio-energetic and development

**DOI:** 10.1101/2020.08.09.243196

**Authors:** János Györi, Andrea B. Kohn, Leonid L. Moroz

## Abstract

ATP and its ionotropic P2X receptors are components of one of the most ancient signaling systems. However, little is known about the distribution and function of purinergic transmission in invertebrates. Here, we cloned, expressed, and pharmacologically characterized P2X receptors in the sea slug *Aplysia californica* – a prominent model in cellular and system neuroscience. We showed that ATP and P2X receptors are essential signaling components within the unique bioenergetic center located in the CNS of *Aplysia*, also known as the cerebral F-cluster of insulin-containing neurons. Functional P2X receptors were successfully expressed in *Xenopus* oocytes to characterize their ATP-dependence (EC_50_=306μM), two-phased kinetics, ion selectivity (Na^+^-dependence), sensitivity to the ATP analog Bz-ATP (~20% compare to ATP) and antagonists (with PPADS as a more potent inhibitor compared to suramin). Next, using RNA-seq, we characterized the expression of P2X receptors across more than a dozen *Aplysia* peripheral tissues and developmental stages. We showed that P2X receptors are predominantly expressed in chemosensory structures and during early cleavage stages. The localization and pharmacology of P2X receptors in *Aplysia* highlight the evolutionary conservation of bioenergetic sensors and chemosensory purinergic transmission across animals. This study also provides a foundation to decipher homeostatic mechanisms in development and neuroendocrine systems.

**Graphical Abstract:** 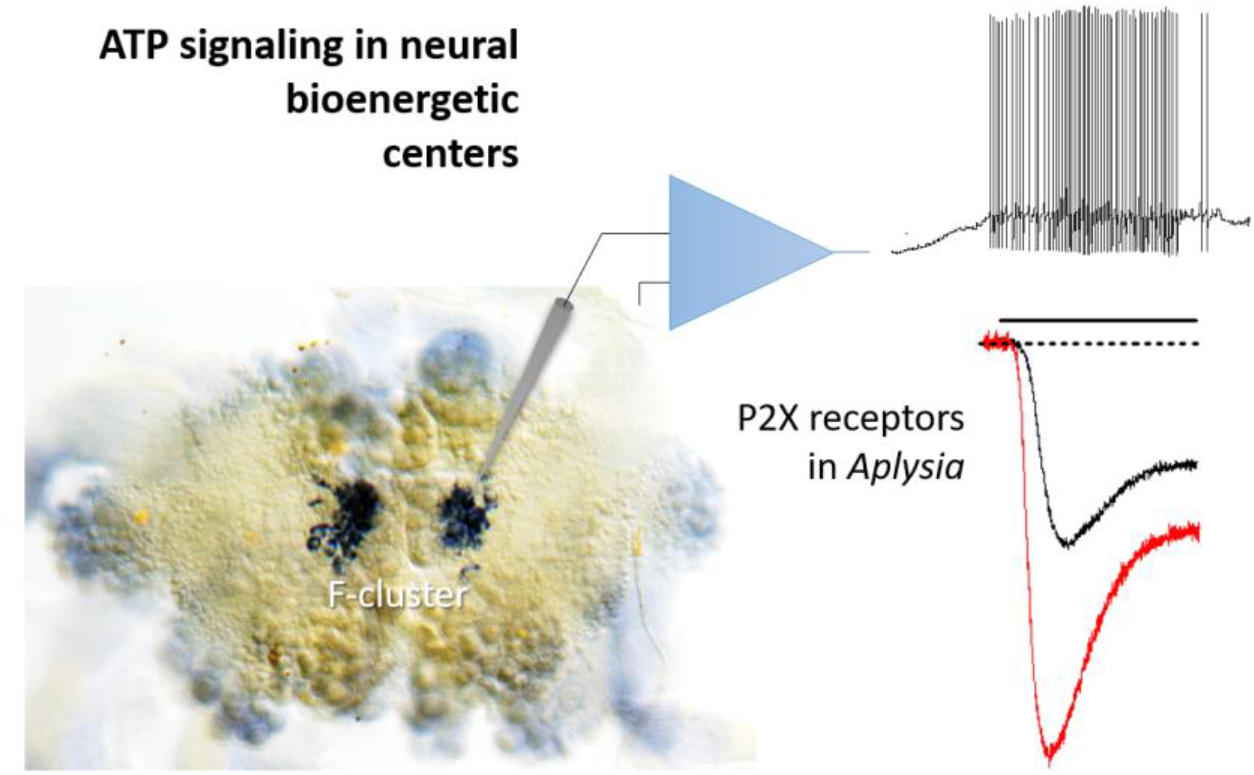

We show that ATP and its ligand-gated P2X receptors are essential signaling components within both the chemosensory systems and the unique bioenergetic center, present in the CNS of the sea slug *Aplysia californica* – a prominent model in neuroscience. Expression and pharmacology of P2X receptors in *Aplysia* confirms the preservation of evolutionary conserved bioenergetic sensors across animals and provide new tools to decipher homeostatic mechanisms in neuro-endocrine systems in general.

## 1. Introduction

In addition to being the critical energy storage for every cell, ATP acts as one of the most ancient intracellular and intercellular signal molecules[1,2]. The possible involvement of ATP in signaling mechanisms was initiated in the 1920s by Drury and Szent-Gyorgyi[3]; and then in the 1950s by Holtons[4–6], leading to the concept of purinergic transmission[7,8]. Eventually, rapid ATP-gated ion currents were discovered in neurons[9,10] and muscles[11], and specific subtypes of the ligand-gated P2X receptors were identified in the 1990s[12–16].

Comparative studies established that P2X-type receptors are broadly distributed across many eukaryotic lineages[1,17–20], including the majority of Metazoa[2,13,21–24]. Nevertheless, the recruitment of P2X receptors into different signaling mechanisms and tissues occurred relatively randomly, and, in some evolutionary lineages, P2X receptors were secondarily lost. For example, *Drosophila* and *C. elegans* genomes have no P2X receptors[25], but other ecdysozoans such as *Daphnia*, the shrimp *Litopenaeus*, the tick *Boophilus*[26] and tardigrades[27] contain one receptor. Many studied Lophotrochosoans, including molluscs and flatworms, also have one type of P2X receptor with shared pharmacological properties to mammals[28]. However, practically nothing is known about the functional roles of P2X receptors in the CNS and peripheral tissues of invertebrates and molluscs, in particular.

The release of ATP from the central ganglia of the pond snail, *Lymnaea stagnalis* was demonstrated[29], and, subsequently, P2X receptors were identified in this species with widespread expression across the CNS[30], but unknown function(s).

Here, we show that ATP and its ligand-gated P2X receptors are essential signaling components within chemosensory structures and the unique bioenergetic center, present in the CNS of the sea slug *Aplysia* – a prominent model in neuroscience[31–36]. Expression and pharmacology of P2X receptors in *Aplysia* confirms the preservation of evolutionary conserved bioenergetic reporter-sensor systems across animals and provide new tools to decipher homeostatic mechanisms in neuroendocrine systems and development.

## 2. Materials and methods

### 2.1. Cloning of Aplysia AcP2X receptors for expression, in situ hybridization and molecular analyses

*Aplysia californica* (60-100g) were obtained from the National Resource for *Aplysia* at the University of Miami (Supplementary Data section#1 for details).

The original sequences were generated using RNA-seq profiling[37–40]. Details for RNA extraction and cDNA library construction have been described[37–39,41] and provided in Supplementary Data. We used the same protocols for whole-mount *in situ* hybridization as reported elsewhere[42,43] with a specific probe for the validated *Ac*P2X (Supplementary Data section#2). Expression of *Ac*P2X was performed in eight experimental and two control preparations of the CNS; additional controls were reported elsewhere[37,43]. Control *in situ* hybridization experiments with full length ‘sense’ probes revealed no specific or selective staining under identical labeling protocols. Images were acquired with a Nikon Coolpix4500 digital camera mounted on an upright Nikon Optiphot-2 microscope.

Expression levels of transcripts were calculated using the normalization method for RNA-seq - Transcripts Per Million (TPM)[44]. Mapping was performed in the STAR (2.3.0)/feature Counts analysis with the values obtained from the Bowtie2/Tophat pipeline[45]. The mapped reads were summarized and counted within the R statistical programming language. Supplementary Methods Section#3 contains a list of SRA RNA-seq projects.

### 2.2. Oocyte expression and electrophysiology

#### RNA and oocyte preparation

RNA was transcribed from full-length cDNAs of *Ac*P2X subunits using the T7 mMessage mMachine *in vitro* transcription kit (Ambion). The amount of purified, transcribed RNA was estimated on a Bioanalyzer (Agilent). Surgically removed stage V and VI oocytes from *Xenopus laevis* were injected with a total of 50ng transcribed RNA (46nL total volume) and incubated at 17°C for three-five days in ND96 medium (96mM NaCl, 2mM KCl, 1mM MgCl2, 1.8mM CaCl2, and 5mM HEPES, pH=7.4) supplemented with 2.5mM sodium pyruvate, 100units/mL penicillin, 100μg/mL streptomycin, and 5% horse serum (Sigma).

#### Oocytes recordings

The oocyte recording bath was in ND96 medium (96mM NaCl, 2mM KCl, 1mM MgCl2, 1.8mM CaCl2, and 5mM HEPES, pH=7.4) with the 1.8mM CaCl2 being replaced by 1.8mM BaCl2. Whole-oocyte currents were recorded by two-electrode voltage clamp (GeneClamp500B, Axon Instruments, Foster City, CA, USA) using microelectrodes made of borosilicate glass (WPI, USA) with a resistance of 0.5–1MΩ when filled with 2.5M KCl. Currents were filtered at 2kHz and digitally sampled at 5kHz with a Digidata 1320B Interface (Axon Instruments, CA). Recording and data analysis were performed using pCLAMP software version 8.2 (Axon Instruments). For data acquisition and clamp protocols, the amplifiers were connected via a Digidata 1320B AD/DA converter (Axon, USA) to an AMD PC with pClamp 8.2 voltage-clamp software (Axon, USA). Unfiltered signals were sampled at 10kHz and stored digitally.

Data are presented as mean+S.E. using Student’s paired *t*-test. Concentration-response data were fitted to the equation I=I_max_/[1+(EC_50_/L)^*nH*^], where I is the actual current for a ligand concentration (L), *n_H_* is the Hill coefficient, Imax is the maximal current and EC_50_ is the concentration of agonist evoking 50% the maximum response. To compute the reversal potential for sodium the Nernst equation used; Vj=(RT)/(zF)ln(c1/c2) where R is the gas constant 1:98 calK^-1^mol^-1^, F is the Faraday constant 96, 840 C/mol, T is the temperature in ^o^K and z is the valence of the ion.

#### *In situ* recordings

Voltage- and current-clamp experiments were carried out on identified F-cluster neurons in intact nervous systems of *Aplysia*[46], ~0.5mL bath was perfused with solutions using a gravity-feed system and a peristaltic pump, and solution exchanges were performed by VC-6 six-channel valve controller (Warner Inst., USA). Conventional two-electrode (3-10MΩ) voltage-clamp techniques (Axoclamp2B, TEVC mode) were employed to measure agonist-activated currents as reported[47] at room temperature(20±2°C). To characterize membrane and action potentials, we used a bridge mode of Axoclamp2B with borosilicate microelectrodes (tip resistance:10-18MΩ, with 0.5M KCl, 2M K-Acetate, and 5mM HEPES, pH=7.2).

## 3. Results

### 3.1 Identity, phylogeny and tissue-specific expression of Aplysia P2X receptors

In *Aplysia*, we identified and cloned a single P2X receptor with two splice forms (GenBank##: NP_001191558.1, NP_001191559.1), which shared 92% identity (Supplementary Data). The predicted structure of the *Aplysia* P2X reveals all major evolutionary conservative sites and posttranslational modifications (Supplementary Data, **Fig. 1S**), which are similar to its homolog in another gastropod, *Lymnaea*[30]. The genomic organization of the P2X receptors confirmed the overall evolutionary conservation of exons and intron-exon boundaries*. Aplysia* P2X receptor exons are similar in number and length to other vertebrate P2X4 receptors, but this is not true in some other invertebrates (Supplementary Data and **Fig. 2S**).

Fig. 1A shows the phylogenetic relationships among P2X receptors with prominent events of gene duplications in the lineages leading to humans, zebrafishes, hemichordates, echinoderms, and basally-branched metazoans such as sponges, placozoans, and cnidarians. In contrast, representatives of molluscs (including *Aplysia*), annelids, parasitic flatworms (*Schistosoma*) seem to have a single copy of P2X-encoded genes, which often form distinct phyletic clusters within a respective phylum. This reconstruction suggests the presence of a single P2X gene in the common metazoan ancestor with independent multiplication events in selected animal lineages. It primarily occurred within vertebrates as the reflection of whole-genome duplications early in the evolution of this group). Interestingly, some bilaterians (such as the acoel *Hofstenia miamia*, insects, and nematodes[48]) secondarily lost P2X receptors. Such mosaic-type phyletic distribution likely illustrates different system constraints for the recruitment of P2X receptors to novel functions and/or preservation of ancestral molecular mechanisms of purinergic signaling.

**Figure 1.**
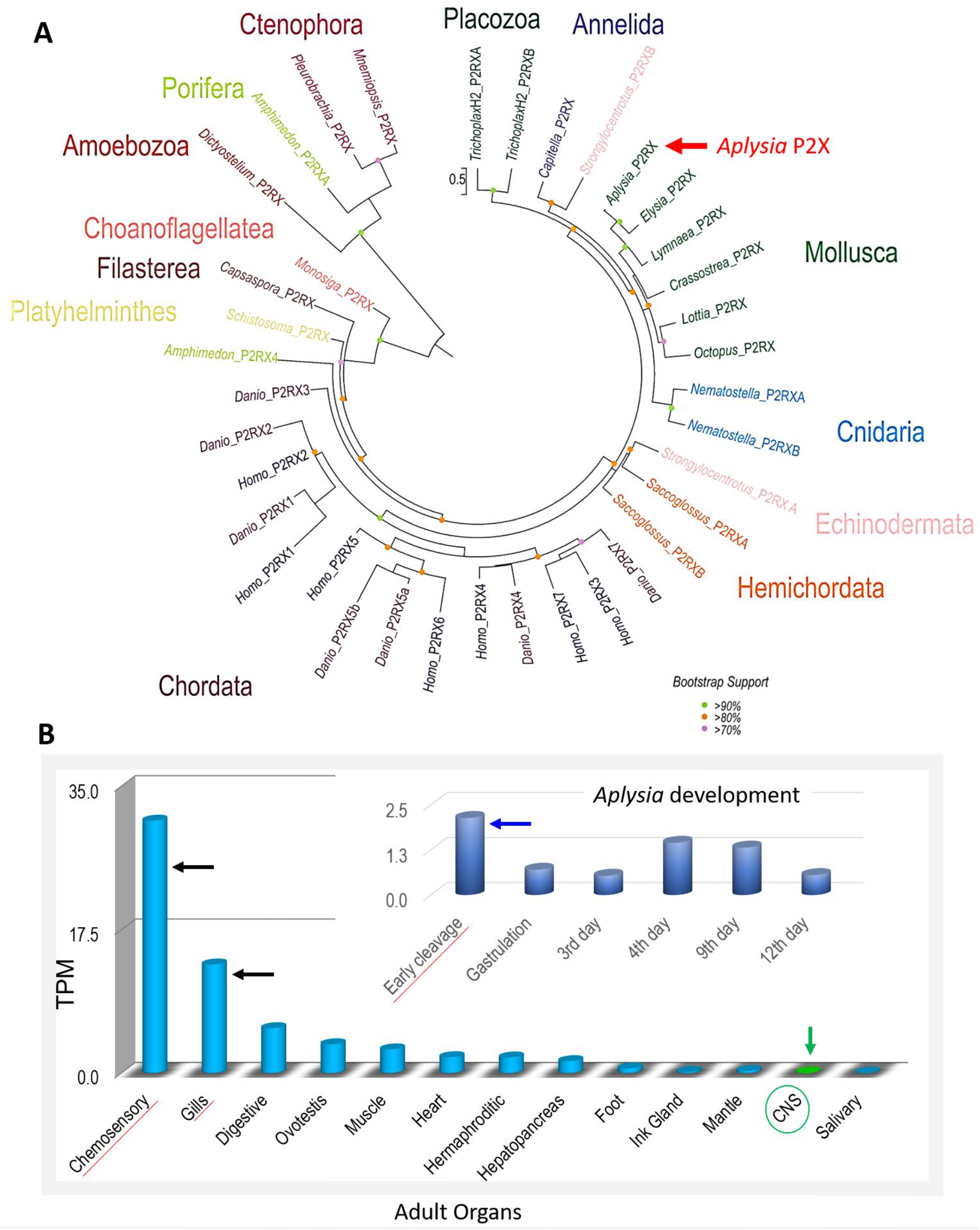
A. **Phylogenetic relationships of P2X and P2X-like receptors (P2RX).** A maximum likelihood (ML) phylogenetic tree of P2X receptors (Supplementary Data#3 and excel table#2 for sequences) with the best-fit model (LG+G). Bootstrap support less than 70 omitted. Phylogenetically, the P2RX predicted proteins cluster by phyla. P2X-type receptors are not unique to metazoans because they are detected in unicellular green algae *Ostreococcus tauri*[18], the amoeba *Dictyostelium discoideum*[17] the unicellular eukaryote *Monosiga brevicollis*[18] as well as *Capsaspora owczarzaki*[19] and all these species appear to have one P2X gene. Most of the non-bilaterians seem to have at least two P2X receptors (except for ctenophores, where only one receptor was detected). Lophotrochozoans, including the mollusc *Aplysia* and kins, appear to have one P2X receptor with different isoforms. The sea urchin and the acorn worm, *Saccoglossus*, both have at least two genes but numerous isoforms[1]. Humans[13], as well as other chordates, appear to have seven unique P2X receptor genes[1]. **B.** Quantification of the expression of P2X receptors in the CNS, peripheral tissues, and developmental stages (insert) of *Aplysia*. The RNA-seq data represented as TPM (<u>t</u>ranscript <u>p</u>er <u>m</u>illion) values[57–59]. The highest expression levels were detected in chemosensory areas (mouth areas and rhinophores), gills, and early developmental stages (Supplementary Data#3 for SRAs).

Next, we characterized the expression of P2X receptors in *Aplysia* using a broad spectrum of RNA-seq data obtained from adult and developmental stages[49] (see Supplementary Data section 3 for details).

The highest level of P2X expression was found in the chemosensory structures (the mouth area and rhinophores[50]) as well as in the gill (**Fig. 1B**), which is also known as the chemosensory and respiratory organ. Expression of P2X receptors was also detected in the majority of peripheral organs of *Aplysia* as well as during the first cleavage stages (**Fig. 1B**), where no neurons or specialized sensory cells exist. Thus, ATP could act as a paracrine messenger in early embryogenesis.

Interestingly, the CNS had the overall lowest level of the P2X gene expression (**Fig. 1B**). This situation might be analogous to the recruitment of purinergic signaling in the chemosensation within the mammalian brain[51], suggesting the presence of a distinct population of ATP-sensing cells. We tested this hypothesis.

*Ac*P2X was explicitly expressed in two symmetrical subpopulations of insulincontaining neurons (Fig. 2A, n=6) localized in the F-cluster in the cerebral ganglion of *Aplysia*[46,52]. Each subpopulation contained about 25-30 of electrically coupled cells (30-50μm diameter, **Fig. 2A-B**)[46]. Application of 2mM ATP to these neurons elicited a 2-5mV depolarization, action potentials, and these effects were reversible (**Fig. 2C**) and voltage-dependent (**Fig. 2D**). Neurons that were negative for *Ac*P2X, by *in situ* hybridization, showed no response to as high as 10mM ATP concentration. These tests confirmed that P2X receptors in F-cluster neurosecretory cells are functional.

**Figure 2.**
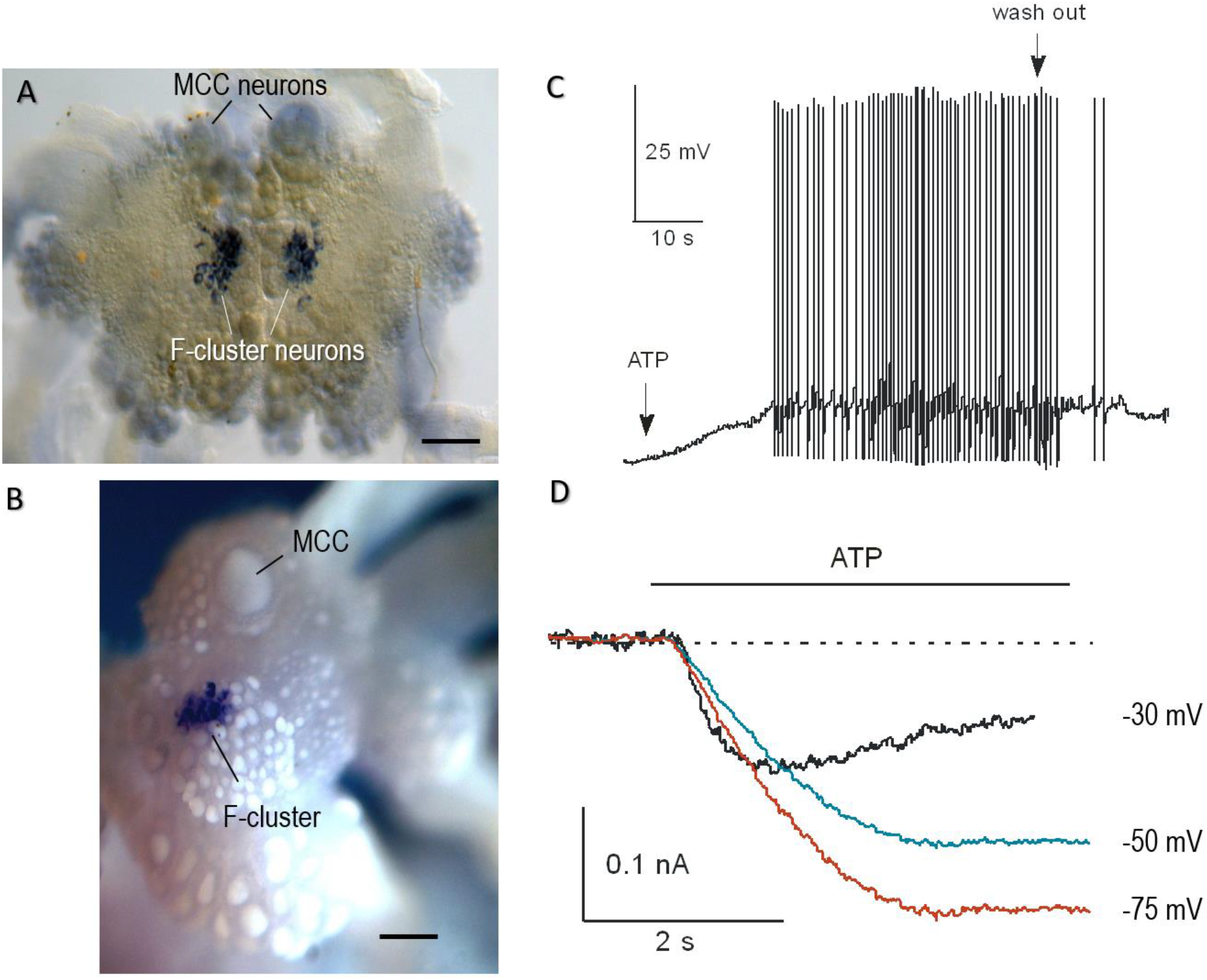
Distribution of *AcP2X* in the CNS of *Aplysia* and the effect of ATP on *Aplysia* F-cluster neurons. A and B: *Ac*P2X is expressed in neurons of the cerebral F-cluster (*in situ* hybridization). A pair of giant serotonergic feeding interneurons (MCC) are indicated by arrows. A. The preparation embedded in a mounting media. B. The cerebral ganglion photographed in 100% ethanol. Scale: 300μm. C: Current-clamp recording from F-cluster neurons in the intact CNS. Bath application of ATP (2.0mM) caused an excitatory response with spiking activity (2-5mV depolarization with a burst of the action potentials), and full recovery following washout (indicated by arrows). D: Voltage-clamp recording from F-cluster neuron. Raw traces recorded in response to 2.0mM of ATP at three holding potentials (agonist application indicated by the line).

### 3.2. Expression of AcP2X in Xenopus oocyte confirms the evolutionary conservation of kinetic and pharmacological parameters

ATP elicited an inward current in a concentration-dependent manner in oocytes injected with *Ac*P2X (**Fig. 3A**). EC50s were determined for both the fast (0-1seconds) and the slow component of current with continuous application of ATP. The EC_50_ for the fast component was 306.0μM with a 1.58 Hill coefficient and for the slow component 497.4μM with a 0.97 Hill coefficient (n=5 oocytes, **Fig. 3B**). The second application of the agonist, with a recovery time of 6 minutes, generated a 15-30% reduction in peak amplitude and is indicative of the rundown observed in other P2X receptor subunits. The response to 250μM ATP produced a mean peak amplitude of 31.3nA+3.8nA and a time to a peak value of 2.76±0.21s (n=19) with a holding membrane potential (HP) of −70 mV (**Fig. 3C**). The ATP analog, 2’,3’-O-(4-B enzoylbenzoyl) adenosine 5’-triphosphate (Bz-ATP[22]) gave a partial response at 20% of the ATP response (n=8 oocyte, **Fig. 3C**). There were no UTP and ADP responses within the same range of concentrations (data not shown).

**Figure 3.**
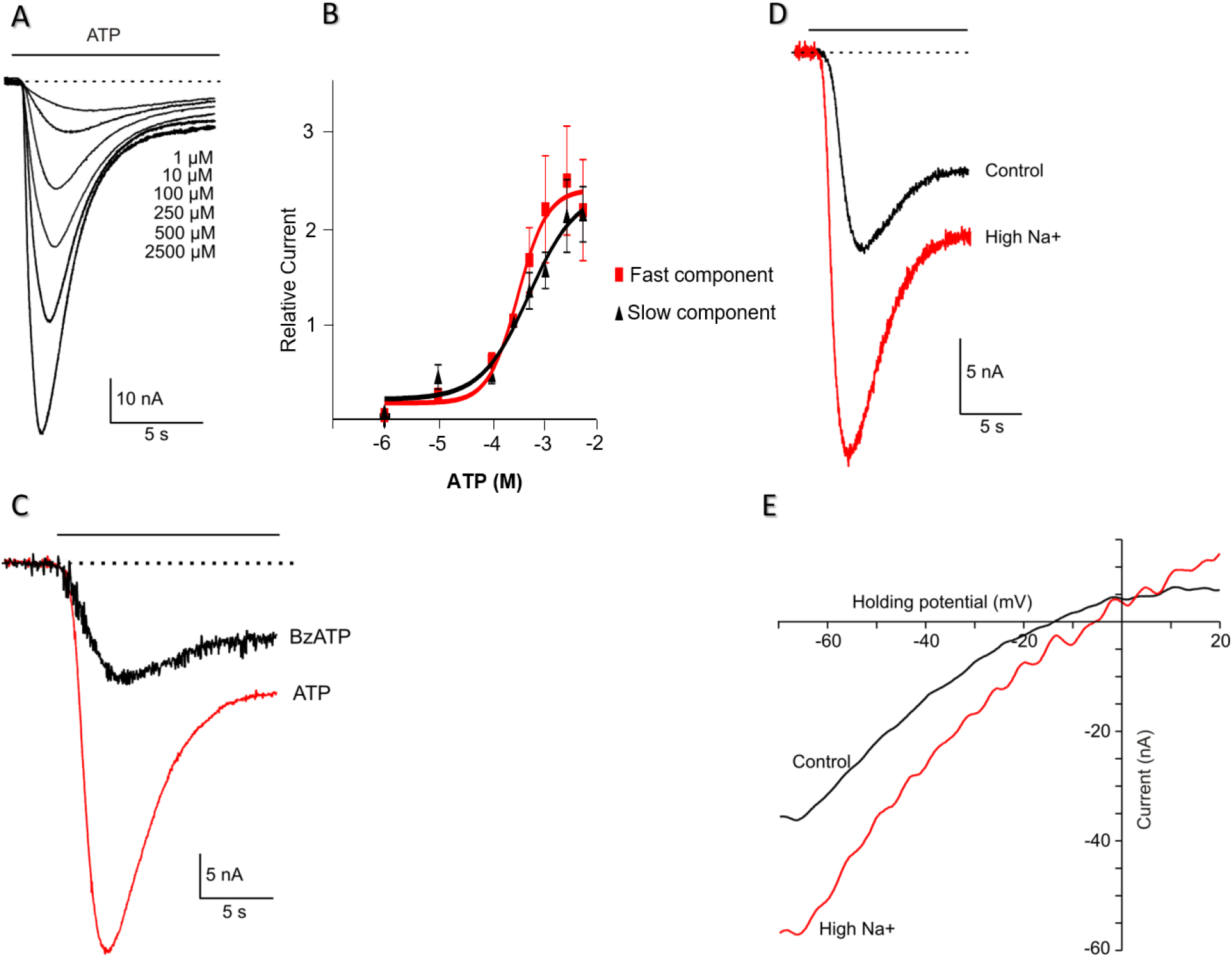
Functional expression of recombinant *AcP2X* receptors in *Xenopus* oocytes. A: Examples of currents recorded in response to different concentrations of ATP (HP=-70mV, agonist application indicated by the solid line). B: Dose-response curves for ATP receptor activation. Mean currents were normalized to the response given by 250μM ATP (*n=*7 oocytes). Serially increasing concentrations of ATP were applied to oocytes for 15s at 6-min intervals. Symbols represent mean±S.E. Continuous line for *ATP* represents data fitted using the equation I= I_max_/[1+(EC50/L)*nH*], where I is the actual current for a ligand concentration (L), *nH* is the Hill coefficient, and I_max_ is the maximal current (EC_50fast_ = 306.0μM, EC_50slow_= 497.4 μM; *nH_fast_=1.58, nH_slow_=0.97*). C. Two-electrode voltage-clamp recordings from oocytes expressing *AcP2X* receptors. Representative inward currents recorded in response to ATP (red trace) and the 250μM of Bz-ATP (HP=-70mV, application indicated by the solid line). D: Recordings of ATP-induced current (250μM, ATP) in the presence of normal [Na^+^] (96mM) and with elevated extracellular Na^+^ (144mM; red trace); HP=-70mV. E: Ramp voltage-clamp protocol from −70mV HP to 20mV in the presence of 250μM ATP. The plots of the subtracted current (a current in the presence of ATP minus the current in the absence of ATP) against voltage during the ramp. The red trace - high [Na^+^], 144mM. According to the Nerst equation, the reversal potential was shifted by 10.2±1.3mV to the + direction of the holding potential.

The current-voltage relationship was investigated in the presence of elevated (144mM) and low extracellular NaCl(96mM) concentrations (n=6 oocytes, **Fig. 3D**). A reversal potential was determined by applying a ramp protocol from −70mV to 20mV in high and normal Na^+^ with 250μM of ATP (**Fig. 3E**). The reversal potential was 13.9mV and shifted by +10.2+1.3mV to positive holding in high sodium solution (n=6 oocytes), according to the Nernst equation.

P2X antagonist suramin[22] inhibited ATP responses in a concentration-dependent manner (**Fig. 4A,B**; 7 oocytes,). Another P2X antagonist, pyridoxal-phosphate-6-azophenyl-2’,4-disulfonic acid(PPADS[22]) also inhibited the response of ATP on *Ac*P2X in a concentration-dependent manner (**Fig. 4C,D**). However, the application of PPADS produced a greater block than the suramin (**Fig. 4E**). Mean current responses to 250μM ATP in the range of 1-250μM PPADS generated an EC_50_=211.2μM for the fast component, but the slow component could not be calculated (5-7 oocytes, **Fig. 4D**). The second splice form of *Ac*P2Xb was also expressed in oocytes producing currents very similar to the first isoform *Ac*P2X described above; however, it resulted in much smaller (and unstable) responses (data not shown).

**Figure 4.**
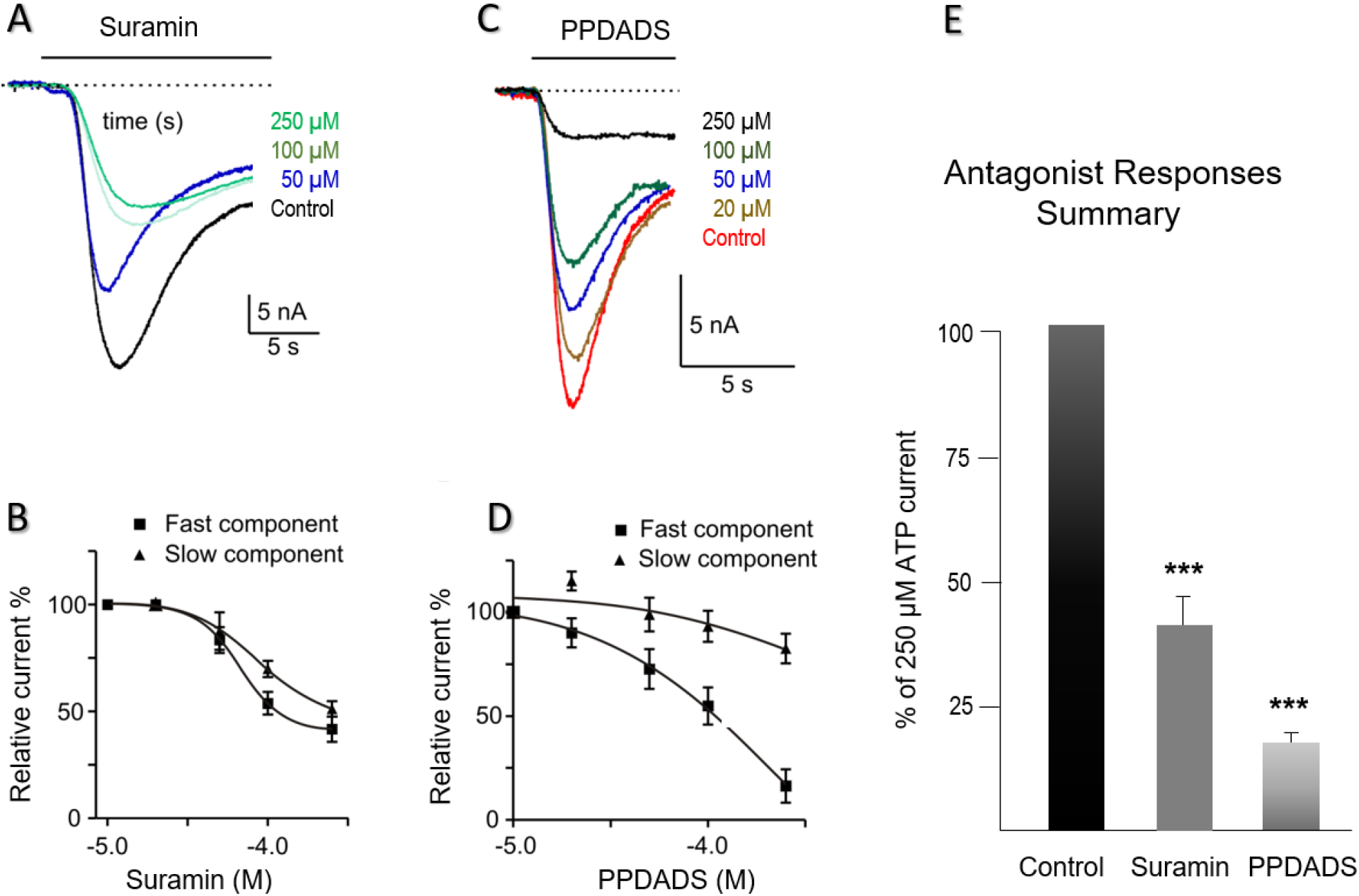
Pharmacology of *Ac*P2X receptors in *Xenopus* oocytes. A: Example of currents induced by 250μM ATP. ATP was applied to oocytes for 15s, in the presence of varying concentrations of suramin (HP=-70mV). B: Mean responses to 250μM ATP in the presence of 1-250μM suramin. There was a suramin-resistant component of the *Ac*P2X current. Symbols represent mean±S.E. C: Traces recorded in response to 250μM ATP in the presence of varying concentrations of the second antagonist, PPADS (concentrations are shown in μM, and all applications are indicated by the solid lines). D: Mean responses to 250μM of ATP in the presence of the PPADS (fast component of responses - closed squares, slow component - triangles). PPADS was an effective antagonist in the range of 1–250μM. Fitting of the data using the sigmoidal dose-response curve by a continuous line, EC_50fast_=211.2. Symbols represent the mean±S.E. E: Among the two antagonists tested, the suramin proved to be a more effective blocker of the ATP-activated channels. A chart of mean currents (% of 250μM ATP response) in the presence of 250μM Suramin and 250μM PPADS. Mean currents were normalized to the response given by 250μM of ATP. Symbols represent mean±S.E; statistically significant differences (Student’s *t*-test) from control (P<0.05) are indicated by asterisks (***) above the bars.

## 4. Discussion

As the central bioenergetic currency, the intracellular concentrations of ATP reach 1-10mM with multiple mechanisms of its extracellular release across all domains of life[2]. In the molluscan CNS, the baseline level of ATP release can be increased following depolarization and application of serotonin, suggesting that ATP can act as an endogenous fast neurotransmitter[29]. The presence of ionotropic P2X ATP-gated cationic channels in peripheral chemosensory structures and the CNS (**Fig. 1B**) further support this hypothesis. However, we also reported P2X receptors early in development, suggesting that ATP might be a paracrine signal molecule controlling cleavage and differentiation.

The purinergic sensory transmission is widespread in mammals[51] and might have deep evolutionary roots[2]. Mammalian P2X receptors[13] are comparable to their homologs in *Aplysia* based on sensitivity to ATP and kinetic parameters. The overall kinetic and pharmacological parameters of *Aplysia* P2X receptors are also similar to those described both in a close-related *Lymnaea*[30] and distantly related *Schistosoma*[28]. However, *Lymnaea* P2X showed much higher sensitivity (EC_50_ is in μmolar range) to ATP, than *Aplysia*. Suramin and PPADS both inhibited the ATP evoked responses in other species[26–28,30], but in *Aplysia*, it occurred in a narrower range (10-250μM) than in *Lymnaea* (0.1μM-250μM).

Ecdysozoan P2X receptors are relatively diverse. In contrast to *Aplysia*, the tick P2X receptor displayed a very slow current kinetics, and little desensitization during ATP application[26]. The tardigrade P2X[27] had a relatively low sensitivity for ATP (EC_50_ ~44.5μM), but fast activation and desensitization kinetics – similar to *Aplysia*.

Thus, *Aplysia* P2X receptors exhibit a distinct phenotype, having a moderate ATP sensitivity (compared to the freshwater *Lymnaea*) but faster kinetics then some ecdysozoans. These “hybrid features” might be related to the marine ecology of *Aplysia* with a broader range of environmental changes.

Interestingly, the abundance of P2X receptors in the *Aplysia* (and kin) chemosensory systems (such as mouth areas, rhinophores, and gills) correlates with the expression of nitric oxide synthase[50,53–55] suggesting interactions of these afferent pathways in the control of feeding and respiration. *Aplysia* might also detect environmental ATP from bacterial and algal (food) sources[2] as in some other studied marine species, including lobsters[56].

Within the CNS of *Aplysia*, P2X receptors are expressed in the distinct cluster of insulin-containing neurons[46,52], likely associated with the systemic control of growth and, subsequently, reproduction. The release of the *Aplysia* insulin can decrease the level of glucose in hemolymph[52].

Moreover, F-cluster neurosecretory cells are electrically coupled[46], which help to synchronize their discharges and, eventually, the secretion of insulin. It is known that ATP can also be released from gap junction (innexins) and during synaptic exocytosis[2]. Thus, we can propose that P2X-induced neuronal depolarization of insulin-containing neurons provide positive purinergic feedback sustaining the excitability and secretory activity of this multifunctional bio-energetic center in *Aplysia* and related gastropods.

## Acknowledgments

We thank Dr. T. Ha and E. Bobkova for help with cloning and in situ hybridization. This work was supported by the Human Frontiers Science Program (RGP0060/2017) and National Science Foundation (1146575,1557923,1548121,1645219) grants to L.L.M.

## Role of authors

All authors had access to the data in the study and take responsibility for the integrity of the data and the accuracy of the data analysis. Research design, writing, acquisition of data: all authors. Molecular data: ABK, LLM; Pharmacological tests: JG; Analysis and interpretation: all authors: Funding: LLM.

